# First Whole-Genome Assembly of the Galápagos Petrel (*Pterodroma phaeopygia*) Using Oxford Nanopore Sequencing to Advance Conservation Genomics in a Critically Endangered Seabird

**DOI:** 10.1101/2025.07.21.665997

**Authors:** Isabella R. Sessi, James B. Henderson, Jessica A. Martin, Alice Skehel, Gabriela Pozo, Vera de Ferran, John P. Dumbacher, Jaime A. Chaves

## Abstract

The Galápagos petrel (*Pterodroma phaeopygia*) is a critically endangered procellariiform seabird endemic to the Galápagos Islands. Once abundant, its populations have sharply declined due to invasive predators, habitat degradation, and destruction of nest burrows. Although the species is distributed across several islands, the demographics of each population and their genetic relationships are poorly understood. To facilitate future studies of population structure and connectivity, we present the first high-quality reference genome for the Galápagos petrel. The genome was assembled from ultra-long Oxford Nanopore sequence data collected from an adult female sampled on San Cristóbal Island. Sequencing was performed at the Galapagos Science Center, enabling local capacity-building and compliance with strict sample import-export regulations of endangered species. The final nuclear genome assembly spans 1.35 Gb in length, with average coverage of 36.07x, scaffold N50 of 74.2 Mb, and a BUSCO aves lineage completeness of 99.95%. The genome comprises 41 pseudo-chromosomes, with 23 spanning from telomere to telomere and 16, including W and Z chromosomes, containing a single telomere. Chromosomal-level scaffolding by reference was performed using the genome of Cory’s shearwater (*Calonectris borealis*) GCA_964196065.2 (Arànega et al., 2024), a related species. The Galápagos petrel reference genome represents a foundational tool for comparative genomics, conservation biology, and functional studies of island-endemic avifauna. It will also facilitate future efforts to characterize genetic diversity, structural variation, and adaptive responses in this critically endangered species.

## Introduction

The Galápagos Islands have long been renowned as a unique biodiversity hotspot, home to thousands of endemic plant and animal species (Kricher & Loughlin, 2022), many of which have small population sizes and rely on a healthy environment to survive (Fernández-Palacios et al., 2021). Since the arrival of humans in the 1500s, hundreds of invasive species have been introduced to the islands, including livestock, rats, cats, dogs, and plants (Causton et al., 2015; Phillips et al., 2012). The introduction of invasive species, compounded with increased human hunting, fishing, and habitat destruction, has reduced native species populations, lowering genetic diversity and pushing some Galápagos endemic species to the brink of extinction (Chaves, 2018; Phillips et al., 2012).

Galápagos petrels (*Pterodroma phaeopygia*), a critically endangered Galápagos endemic, have a large range outside the archipelago when not breeding, extending across the eastern Pacific from the coasts of Mexico to Peru and westward in the tropics to French Polynesia (BirdLife International, 2018; Granizo, 2002). When breeding, Galápagos petrels return to nesting colonies on five islands: Santa Cruz, San Cristóbal, Floreana, Santiago, and Isabela (Cruz-Delgado et al., 2010). Each of these populations is demographically isolated from the others due to nest site fidelity; that is, individual petrels born on a given island return to the same island to breed each year (Cruz & Cruz, 1990).

Galápagos petrels face ongoing population declines from human-induced threats (Zurita-Arthos et al., 2023). As a burrowing species, they are particularly sensitive to invasive plants that grow over nest entrances and make nesting territories inhospitable (Zurita-Arthos et al., 2023). They fall prey to invasive rats, cats, dogs, and feral pigs, which raid nests, predating on eggs, young (Tapia-Jaramillo et al., 2025), and adult petrels (Duffy, 1984; Harris, 1970). More than 90% of the petrels’ nesting habitat is located on privately owned land, leading to increased habitat loss due to agricultural development (Cruz-Delgado et al., 2010; Zurita-Arthos et al., 2023). Grazing livestock in such areas trample and ruin petrel nesting areas (Cruz-Delgado et al., 2010; Duffy, 1984). These anthropogenic factors have led to low breeding success, and Galápagos petrel populations have declined dramatically, leading to their listing as a critically endangered species by the IUCN (BirdLife International, 2018; Cruz-Delgado et al., 2010; Tomkins, 1985).

Previous studies on the genetics of the Galápagos petrel have been limited in scope and primarily focused on a few microsatellites, leaving much unknown about the genetic status of this critically endangered species (Browne et al., 1997; Friesen et al., 2006). A whole-genome assembly for this species would enable detailed population genomics studies, key tools for assessment, monitoring, and management of the species (Brito & Edwards, 2009).

Recent studies have highlighted the importance of genomic data in advancing conservation efforts (Hogg, 2024; Romanov et al., 2009; Theissinger et al., 2023), but these often depend on access to substantial technological infrastructure. The improvement of low-cost portable sequencing platforms offers a solution that is well-suited for use in remote and logistically challenging environments (Watsa et al., 2020) such as the Galápagos Islands. Sequencing on the island served two key purposes: (1) building local capacity and supporting local conservation genomics efforts, and (2) abiding by the regulations and control for samples of species protected under the United States Endangered Species Act (ESA) and the Ecuadorian implementation of the Nagoya Protocol (Secretariat of the Convention on Biological Diversity, 2010).

Here we present the first assembled whole-genome of the critically endangered Galápagos petrel (*Pterodroma phaeopigia*), sequenced in the Galápagos, including both W and Z sex avian chromosomes.

## Methods and materials

### Specimen collection

Approximately 50 µl of blood was drawn from the brachial vein of an adult female Galápagos petrel, using a 27-gauge needle and a 1ml syringe on the 2nd of July 2024 in La Comuna, San Cristóbal Island (0°53’04.5"S 89°27’54.9"W). Sample collection was conducted in accordance with the regulations of the authorities and guidelines for the use of birds in field research. The sample was preserved in NAP buffer (Camacho-Sanchez et al., 2019) at the collection site and stored at -80 °C at the Galapagos Science Center (GSC) until DNA extraction.

### DNA isolation and sequencing

High molecular weight (HMW) DNA was extracted using the Monarch UHMW DNA Extraction Kit for Cells and Blood (NEB #T3010, New England Biolabs, Ipswich, MA) following the manufacturer’s protocol. DNA concentration was measured using Qubit Fluorometric Quantitation (Thermo Fisher Scientific). A library was prepared with the Oxford Nanopore Technologies (ONT) Ultra-Long DNA Sequencing Kit (SQK-ULK114), and sequenced on three PromethION 2 flow cells (FLO-PRO114M) using the PromethION 2 Solo (P2 Solo) platform. A second library was prepared from the same sample in January 2025 during a follow-up field trip, using the same protocols and sequenced using two flow cells.

Both sequencing runs were conducted on-site at the GSC using a portable Windows DELL laptop (see Supplement: DELL Laptop Specs) connected to the P2 Solo device. Raw sequence signal was saved to the laptop in POD5 files, a streaming format for direct sequencing instrument output (github.com/nanoporetech/pod5-file-format). ONT was selected due to its portable sequencing capabilities and ability to generate ultra-long reads for assembling through complex and repetitive genomic regions with newer R10 flow-cell Kit 14 chemistry and improved software (github.com/nanoporetech/dorado) (Cheng et al., 2025; Stanojević et al., 2024), producing reads with error profiles suitable for current long-read assembly programs.

### Basecalling, adapter-trimming, and error-correction

Each of the two runs was independently basecalled and adapter-trimmed as described below. The two resulting trimmed-only files were concatenated into a single fastq and used as input to Hifiasm v0.24 (Cheng et al., 2021, 2025), which was released during the preparation of this assembly. This version of Hifiasm performs its own error correction for ONT simplex R10 reads. The resulting output created the first assembly.

Separately, the trimmed reads from each run were also error-corrected using Dorado v0.9.1 (see below). These two Dorado corrected read sets were used as input for the Flye assembler and for a previous version of Hifiasm (v0.20). The resulting output created the second and third assemblies (Figure S3).

POD5 files, copied from the laptop computer used during sequencing at the GSC, were basecalled on a GPU-enabled computer using Dorado v0.7.4 basecall in super-accurate mode (SUP) via MinKNOW Core v6.0.8. This produced multiple 4K fastq files, which were combined into complete datasets for each run.

Adapters were trimmed using porechop_abi v0.5.1 (Bonenfant et al., 2022) with the --ab_initio argument. Trimmed reads were then error corrected with Dorado v0.9.1 using arguments correct –cpu -t 96 on an HPC Linux server, producing fasta-formatted corrected reads.

Read statistics and quality metrics for trimmed-only reads, Hifiasm corrected reads, as well as a subset of metrics for Dorado corrected reads, were calculated using NanoPlot (De Coster et al., 2018) and a custom script (seq_summary_qscore_lens.sh, see Code Availability).

### Genome assembly and scaffolding

Initial read coverage, genome size, and heterozygosity levels were estimated using GeneScopeFK (https://github.com/thegenemyers/GENESCOPE.FK), with kmer sizes 21, 25, and 27. This is a command line implementation of GenomeScope2 (Ranallo-Benavidez et al., 2020) with identical results, built-in kmer histogram production, and plot figure file creation.

Three assemblies were produced for comparison: (1) Hifiasm v0.24, which used trimmed reads with the –ont option, (2) Flye v2.9.5, which used error-corrected reads input as –nano-corr along with the –scaffold option; and (3) Hifiasm v0.20, run with the same corrected reads used for Flye.

Resulting assemblies were checked for adapter and other contamination using the NCBI Foreign Contamination Screen tools (https://github.com/ncbi/fcs), FCS-adaptor, and FCS-GX, against taxonomy ID 53680, and evaluated for standard metrics including N50, L50, using in-house modified script asmstats.pl (see Code Availability) and for genome completeness via Compleasm v0.2.6 (Huang & Li, 2023), miniprot (Li, 2023), and BUSCO v5.4.7 (Simão et al., 2015), MetaEuk (Karin et al., 2020), with the 8338 ortholog aves_odb10 lineage database. Statistics from asmstats, seqtk telo (github.com/lh3/seqtk), and combined Compleasm, BUSCO output were compiled after major pipeline stages to assess modifications. Assembly statistics (Table 1) led us to select Hifiasm v0.24 as the assembler with which to proceed.

**Table 1.** Comparison of the three contig-level assemblies for *Pterodroma phaeopygia*. Bolded values indicate the best or tied for the best category; italics indicate the second best.

| Metric | Hifiasm v0.24 | Flye v2.9.5 | Hifiasm v0.20 |
| --- | --- | --- | --- |
| Number of contigs | **109** | 1038 | *918* |
| Total contig length | 1.408G | 1.31G | 1.446G |
| Longest contig | **207.299M** | *104.336M* | 26.217M |
| N50 / L50 | **36.27M / 8** | *20.93M / 17* | 7.29M / 64 |
| N90 / L90 | **8.79M / 39** | *2.14M / 93* | 1.17 / 234 |
| auN50 | **69.18M** | *32.81M* | 8.19M |
| BUSCO counts, n:8338 | **C:8334, D:58, F:3, M:1** | *C:8333, D:58, F:4, M:1* | C:8329, D:623, F:7, M:2 |
| BUSCO percentages, | **C:99.95%, F:0.04%,** | C:99.94%, F:0.05%, | C:99.89%, F:0.08%, |
| n:8338 | **M:0.01%** | **M:0.01%** | M:0.02% |

Contaminants, primarily mitochondrial, with minimal viral or bacterial presence, were identified and filtered by assembling the mitochondrial genome (see Mitochondrion assembly and annotation section) and comparing the resultant complete mitochondrion against the nuclear genome via blastn (Altschul et al., 1990; Camacho et al., 2009). To avoid removing nuclear-integrated mitochondrial sequences (NUMTs), a minimum 80% read length mapping was required to identify a read as a mitochondrial contaminant.

After removing reads flagged as mitochondrial or foreign contamination, the selected assembler, Hifiasm v0.24, was run with this cleaned input to generate a consensus primary contig-level genome. Assembly output is in gfa format and not the fasta format required by downstream processing (gfa-spec.github.io/GFA-spec), so we converted the gfa to fasta and extracted circular contigs to a separate file with a custom script using awk (see Code Availability). The purge_dups v1.2.5 (Guan et al., 2020) pipeline was run to identify what it terms Haplotig, Repeat, and Junk elements in the contig assembly, which it removes to create a new assembly fasta file. Additionally, two contigs identified in BUSCO analysis as duplicates were excluded.

We performed reference-based scaffolding (Alvarez-Costes et al., 2025; Dunn, 2024; Jorquera et al., 2025) with RagTag v2.28 (Alonge et al., 2022) to provide contig groupings, locations, orientations, and confidence intervals. As a scaffolding reference, we used the NCBI genome GCA_964195595.2 Cory’s shearwater (*Calonectris borealis*), which was chosen as the closest procellariiform relative with a high-quality chromosome-level genome assembly. The hap1 assembly, not NCBI RefSeq hap2, was chosen since hap1 includes W, Z, as well as the 39 numerically identified autosomal pseudo-chromosomes. Single copy gene presence/absence was cross-validated against the Cory’s shearwater information using BUSCO Aves lineage orthologs. Gap filling of the scaffolded assembly was conducted using quarTeT GapFiller v1.2.1 (Lin et al., 2023) with the reference scaffolded assembly and Hifiasm corrected reads as input. Records in homology with reference pseudo-chromosomes have names prefixed *Pphae* and numbered as the *C. borealis* homolog; the rest were prefixed *unloc* to indicate unlocalized sequence to any of the *C. borealis* pseudo-chromosomes (bPteroPhaeo_1.0.fasta).

### Repeat analysis

Before gene annotation, *de novo* repeats were identified using RepeatModeler v2.0.1 (Flynn et al., 2020), which uses RECON (Bao & Eddy, 2002), RepeatScout (Price et al., 2005), and LtrHarvest/Ltr_retriever (Ellinghaus et al., 2008; Ou & Jiang, 2018). Repeat models identified through *de novo* analysis of the assembly were combined with curated Aves models from Dfam 3.8 (dfam.org/releases/Dfam_3.8) and used as input for RepeatMasker v4.0.9 (Smit et al., 2013–2015). This process generated a table summarizing repeat types and lengths, as well as a repeat-masked (soft-masked) version of the assembly, which was subsequently used for gene model annotation.

### Genome annotation

We performed *ab initio* gene model annotation using BRAKER3 v3.0.8 (Gabriel et al. 2023) run in EP mode (clade-level database of protein families for modeling, no species RNAseq) with the soft-masked genome assembly and GeneMark-EP+ and AUGUSTUS employing the vertebrate protein database from OrthoDB v11 (Brůna et al. 2021; Hoff et al. 2019; Stanke et al. 2008, 2006). We added argument --busco_lineage=aves so BRAKER3 invokes compleasm with this lineage and refines results with its output. The gff3 annotation, coding sequence DNA, and protein sequence amino acid files were then used as input for further refinement and functional annotation.

Gene models without start and stop codons were removed from the BRAKER3 output files, as were models contained completely within another. Protein domains were identified in the amino acid (AA) sequences by running InterProScan v5.72-103.0 (Jones et al. 2014). The sequences, DNA or AA as appropriate, were also searched against GenBank databases (downloaded Jan 11, 2025) using blastn with nt database, blastp with uniprot_sprot, and diamond blastp (Buchfink et al. 2021) with TrEMBL and nr databases. Diamond blastp was also used to search OrthoDB v11 vertebrate protein AA sequences. Sequences were additionally aligned to the eggNOG v5.0.2 database with eggNOG-mapper v2.1.11 (Cantalapiedra et al. 2021). Protein domain IDs and Gene Ontology terms from the InterProScan output were added to the gff3 and sequence files for each gene model, as was the functional annotation description from the lowest eValue for each gene from the blast or eggNOG searches when one or more were found with eValue lower than 1e-10.

Gene models were evaluated with BUSCO in protein mode, lineage aves_odb10, and with OMArk v2.0.3 (Nevers et al., 2025) using the OMA LUCA.h5 database to assess proteome completeness with Hierarchical Orthologous Groups (HOGs). Statistics were generated from the annotation-augmented gff3 file with a custom script (basic_gff_stats.sh, see Code Availability).

### Mitochondrial assembly and annotation

HiFiMiTie v0.08 (github.com/calacademy-research/HiFiMiTie), a long-read-based program for mitochondrial assembly, was used with reads previously corrected in a preliminary Hifiasm v0.24 run. This approach facilitated both the removal of mitochondrial contaminants from the nuclear genome assembly input and the generation of a complete mitochondrial genome for this species. Reads matching Aves entries in the NCBI mitochondrial database (downloaded Jan 11, 2025) with 50% or greater sequence coverage from a blast search were used to construct a consensus mitochondrion, its annotation, and analysis of the *control region* heteroplasmy. The Aves taxonomy was also used to define the canonical starting tRNA for the sequence and to select the appropriate mitochondrial genetic code. MITOS2 (Donath et al. 2019; Bernt et al. 2013), running locally, was used to confirm results produced by the pipeline. A circular mitochondrial depiction was created with Geneious Prime 2025.1.2 (www.geneious.com) by importing the fasta sequence and gff annotation, setting the sequence as circular, and then saving the displayed figure with the File > Save as image file option.

## Results

### Basecalling, filtering, and correction

The two basecalled runs produced 4,101,873 reads totaling 49.1 Gbp. Trimmed reads have 4,103,512 reads, slightly more due to reads split by porechop_abi, and 48.69 Gbp with an average coverage for a 1.35 Gbp genome of 36.07x, mean read length 11,866, and N50 22,496. Almost all reads have average quality Q10 or greater, and those with Q20 or greater have over 23x of coverage (Table S1).

Dorado corrected reads reduced the read count to 2,403,479, though the number of bases retained, at 42.66 Gb, is 87.6% of the pre-corrected bases, representing 31.6x genome coverage. N50 improved slightly to 24,321. Quality scores are not available since Dorado correct output is in fasta file format, however, the method is intended to raise the read mean quality to a minimum of Q10. Though relative to Hifiasm –ont, “on a few limited datasets herro [i.e. Dorado correct] gives slightly better QV (by ∼1 in the Phred scale)" (github.com/chhylp123/hifiasm/issues/742).

Hifiasm corrected reads increased Q20 mean read bases by 25% (39.04 Gbp after, 31.16 Gbp before) and more than 3 1⁄2 times more Q25 mean read bases (18.68 Gbp after, 5.1 Gbp before).

### Genome size estimate

Genome size estimates of the 21, 25, and 27 length kmer histograms ranged from 1.238 Gb to 1.239 Gb, heterozygosity 0.47% to 0.54%, and repeat length of 8.95% to 10.57% of the size, a likely underestimate of the repeats, as results following indicate. Unique length estimates ranged from 1.108 Gb to 1.128 Gb, with the final assembly’s non-repeat length in the middle of this range.

### Genome assembly

The Hifiasm v0.24 –ont contig assembly scored best in six metrics used to assess assembly quality (Table 1). Hifiasm v0.24 uses its own error correction, whereas Hifiasm v0.20 used dorado-corrected reads, as did Flye. The Flye v2.9.5 metrics were second in each category except the number of contigs. Compleasm plus BUSCO aves_odb10 results were near full completeness for all three, with a single BUSCO missing in each of the top two assemblies (OrthoDB 48114at8782) and two missing in the other (Table 1, Tables S2-S4).

These assembly results contrast with those found in the assembly of the lava gull (*Leucophaeus fuliginosus*), in which Flye was the top-performing assembler (Martin et al., 2025). These contrasting results are notable given that both studies used the same laboratory and sequencing protocols; however, our study included two sequencing runs, resulting in approximately 60% greater coverage. Thus, the performance difference in our assemblies may reflect Hifiasm’s ability to take full advantage of increased depth through its built-in error correction and graph-based haplotype resolution, both of which benefit significantly from higher coverage.

Hifiasm v0.24 was rerun after removing contaminant-identified reads, resulting in identical output, confirming FCS results finding no contamination in the assembly, and that any mitochondria had been assembled in circular sequences, which were excluded in gfa to fasta conversion. Purged_dups yielded an assembly of 1.346 Gb with 61 contigs and reduced BUSCO duplicates to 31, increased N90 to 10.8M from 8.8M, and left the largest contig, N50, and the single Missing BUSCO unchanged (Table S10).

The RagTag scaffolded genome with *C. borealis* hap1 used as a reference contains 44 records after gap filling. Twelve records were scaffolded with two or three input contigs (eight 2, four 3 contigs), introducing 16 gaps, the rest remaining as contigs. Four of these 16 gaps were filled in three scaffolds, converting them to contigs with no gaps present. This includes the largest in the assembly, pseudo-chromosome 1, which also has both telomeres. Table 2 shows the progression from the initial contig assembly, the duplicate purged assembly, and finally the gap-filled scaffolded assembly containing 12 gaps we have named bPteroPhaeo_1.0. (Table 2).

**Table 2.**
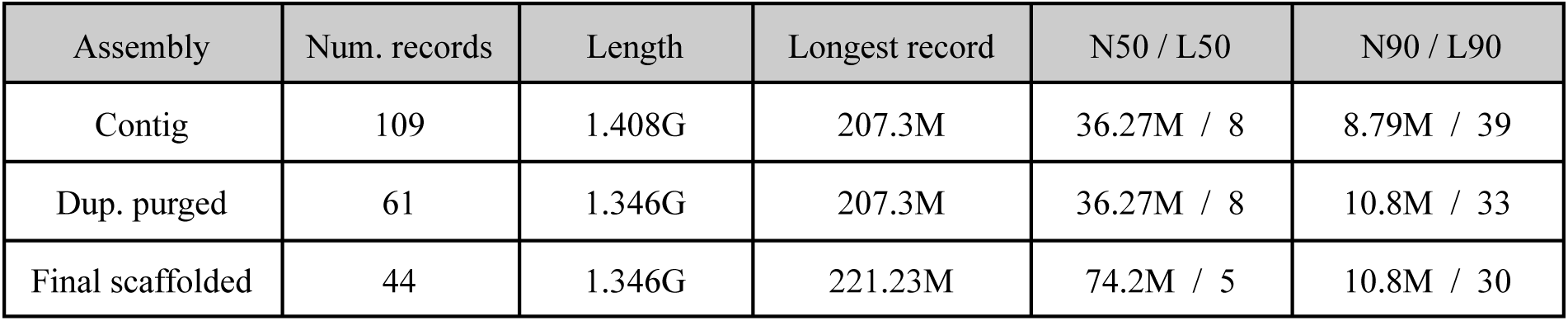
Improvements through the three stages of the assembly pipeline.

| Assembly | Num. records | Length | Longest record | N50 / L50 | N90 / L90 |
| --- | --- | --- | --- | --- | --- |
| Contig | 109 | 1.408G | 207.3M | 36.27M / 8 | 8.79M / 39 |
| Dup. purged | 61 | 1.346G | 207.3M | 36.27M / 8 | 10.8M / 33 |
| Final scaffolded | 44 | 1.346G | 221.23M | 74.2M / 5 | 10.8M / 30 |

The final genome assembly is approximately 1.35 Gb in size, consistent with, though on the upper end of, medium-sized seabird genomes (e.g., *C. borealis* at 1.36 Gb, 1.2 Gb *Pelecanoides urinatrix* (NCBI GCA_013400755.1)). A scaffold N50 of 74.2 Mb and L50 of 5 indicate that half of the genome assembly is contained within just five scaffolds, consistent with the largest macrochromosomes typically observed in avian karyotypes (Kiazim et al., 2021; Kretschmer et al., 2018). In homology with *C. borealis,* we identified 41 records as pseudo-chromosomes, of which 23 are T2T and 16 have one telomere, including the sex chromosomes (i.e., W and Z) (Table S5). All but one of the 8,338 BUSCO aves_odb10 orthologs are found. These values indicate a high-quality genome assembly (Figure 2A, Table S6).

**Figure 1.**
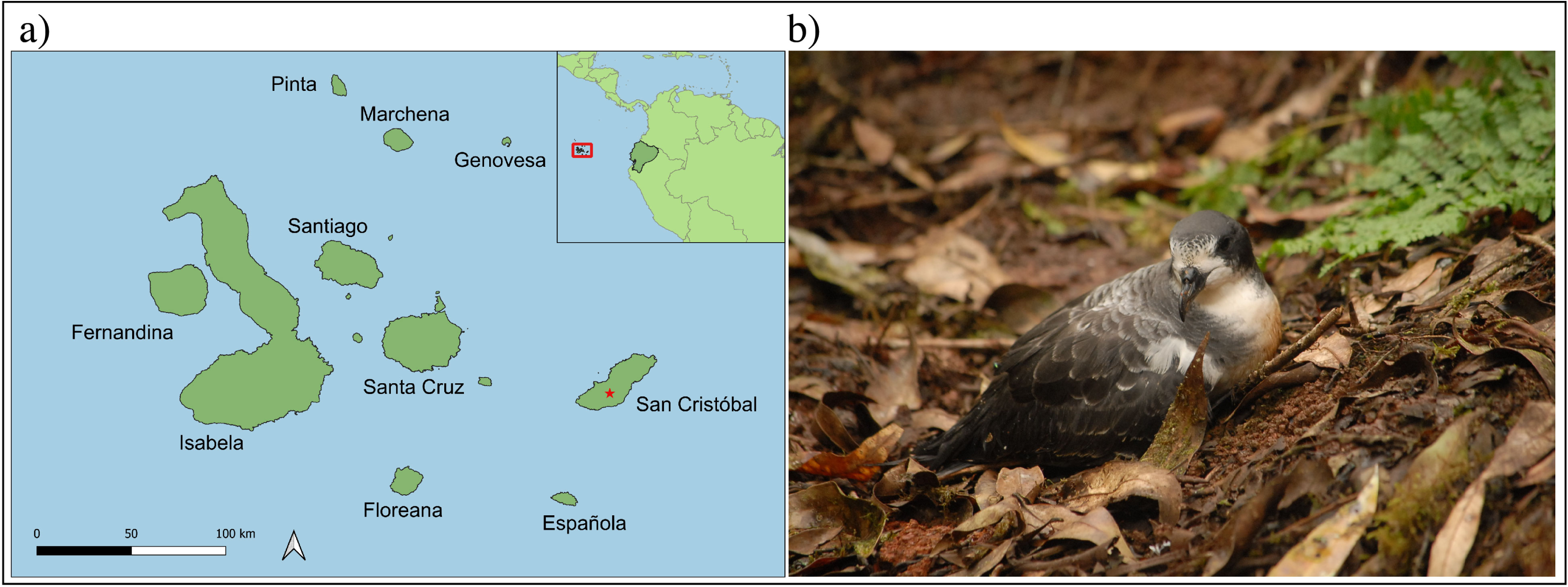
Galápagos petrel and Galápagos Islands. **a)** Sample location at La Comuna, San Cristóbal Island is shown with a red star. **b)** A Galápagos petrel outside its burrow in the highlands of San Cristóbal (Photo: J. Chaves).

**Figure 2.**
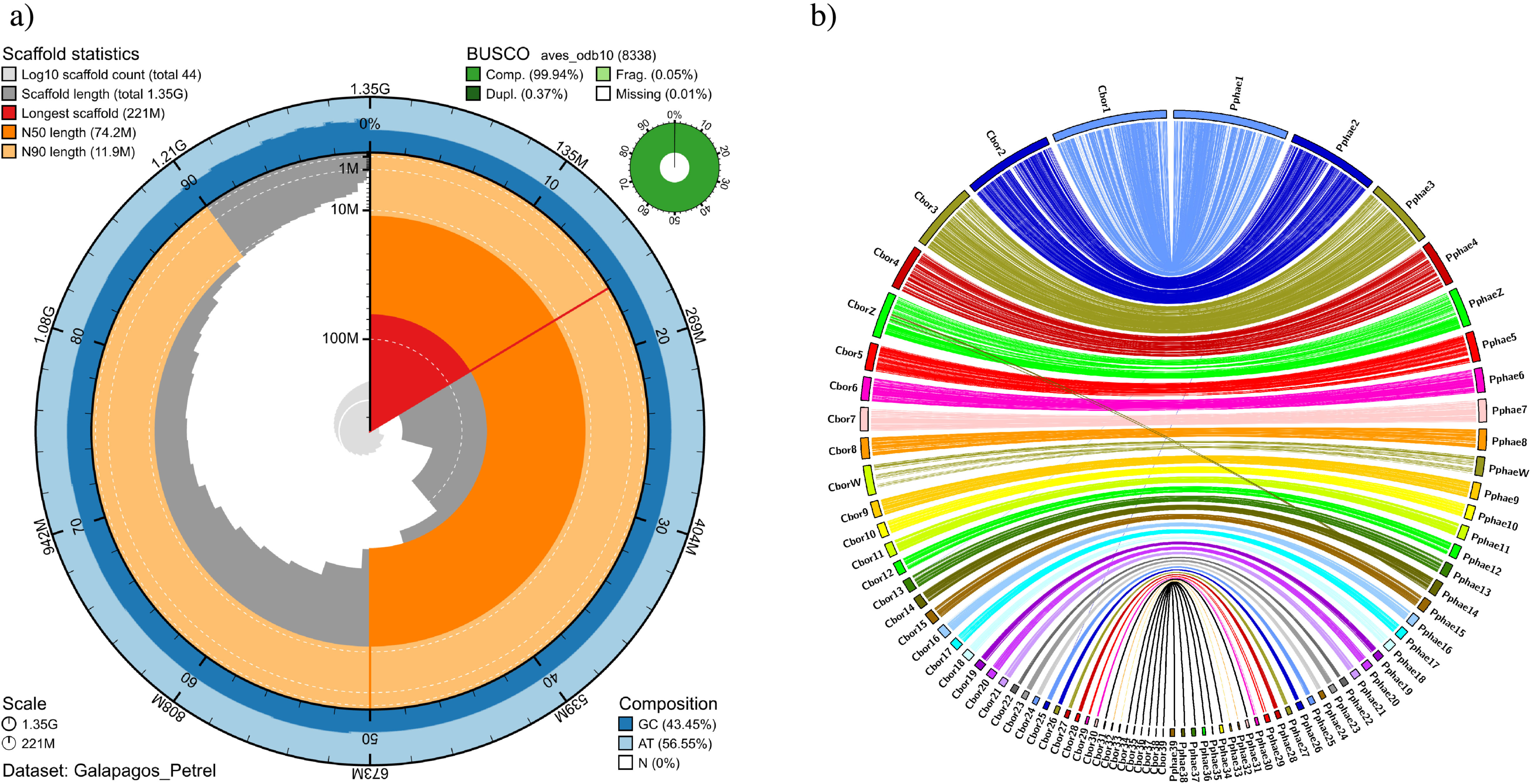
Snail plot of Galápagos petrel and synteny between Galápagos petrel and Cory’s shearwater. **a)** Blobtoolkit snail plot presentation of assembly metrics. **b)** Synteny between *P. phaeopygia* and *C. borealis* using Circos plot.

The level of synteny between the *P. phaeopygia* and *C. borealis* assemblies based on BUSCO orthologs from the aves_odb10 dataset identified only one missing BUSCO for this assembly (*bPteroPhaeo_1.0*), while the *C. borealis* assembly is missing that same ortholog plus one additional gene (Figure 2B). As a result, nearly all of the 8,338 aves_odb10 BUSCOs are represented in the comparison, underscoring the high genomic completeness and structural similarity between the two assemblies. Several microchromosomes lacked annotated BUSCOs in both assemblies. For these regions, we used RagTag alignment results to identify sequence agreement and visualized them as black ribbons in the Circos plot (Figure 2B).

### Repeat analysis

RepeatMasker analysis using the Dfam28 curated Aves dataset combined with *de novo* repeats from RepeatModeler reports 17.43% of the total assembly sequence as repeats at 234.7 Mb. The 1.112 Gb total non-repeat sequence is 3 Mb from the kmer 21 GenomeScope2 estimate, less than a 1% difference; whereas, the repeat content estimates are each at least 100 Mb less than the repeat analysis finds. (Figure S4: Repeat classification, Table S8: Chromosomal and unlocalized record repeat percentages).

### Genome Annotation

BRAKER3 *ab initio* annotation, after refinement requiring start and stop codons and excluding embedded genes, presented candidate models for 29,316 genes and 31,448 mRNAs (genes and isoforms). Functional analysis, which requires identified protein domains or homology with existing genes, retained models for 27,344 genes and 29,343 mRNAs. Compleasm running in protein mode found 7811, 93.68%, of the aves BUSCOs in the protein AA sequences (Table 3). OMArk identified input as belonging to Neognathae and reported results from the Hierarchical Orthologous Groups associated with this clade (Table 4, Figure S8).

**Table 3.** Basic gene model statistics.

| Annotation type | Value |
| --- | --- |
| Number of genes | 27,244 |
| Genome percentage | 15.23% |
| Number of mRNA | 29,343 |
| mRNA mean length | 8,143 bp |
| Mean exons per mRNA | 6.56 |
| Mean mRNA exon length | 174.46 |
| Single exon genes | 5,895 |
| Named genes | 20,130 |
| Named mRNA | 22,188 |
| BUSCO aves_odb10 lineage | 93.68% / 7811 |

**Table 4.**
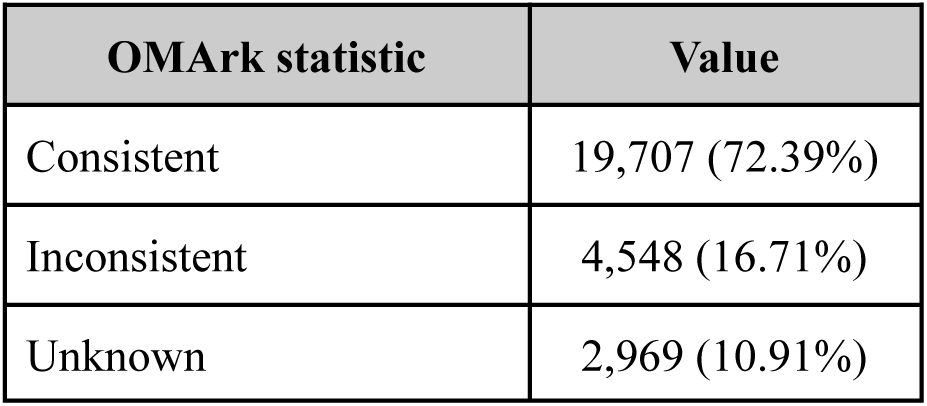
Select OMArk Neognathae HOGs statistics.

### Mitochondrial assembly and annotation

The *P. phaeopygia* consensus mitochondrial sequence derived from 324 mitochondrial HiFi reads is 17,315 bp with 22 tRNAs, 13 protein coding genes, and two rRNAs, as typical in birds and most vertebrates (Table S9, Figure S7). The control region of 1,748 bp has a 138 bp consensus repeat motif repeated 3.9 times spanning 538 bp. This small number of mitochondrial records is likely due to the use of the ultra-long method with selected molecules for sequencing typically longer than vertebrate mitochondria.

## Discussion

The successful generation of a high-quality reference genome for the critically endangered Galápagos petrel (*Pterodroma phaeopygia*) represents a significant contribution to seabird genomic research. Our assembly, generated from ultra-long Oxford Nanopore Technologies (ONT) reads and scaffolded using a closely related species, demonstrates exceptional quality with a scaffold N50 of 74.2 Mb and only 12 gap regions. It achieves near-complete gene content (i.e., BUSCO aves_odb10 score of 99.94% and only a single missing ortholog) and chromosomal-level resolution (i.e., 41 pseudo-chromosomes: 39 autosomal, W and Z, including 23 telomere-to-telomere). These metrics are consistent with benchmarks for high-quality reference genomes (Huang & Li, 2023; Park et al., 2023), supporting its utility in downstream applications.

This assembly is the first genome published for a species in the genus *Pterodroma* and only the second within the Procellariidae family at chromosomal resolution, filling a significant taxonomic gap. Previous genetic research on the Galápagos petrel was limited to microsatellite markers and mitochondrial DNA (Browne et al., 1997; Friesen et al., 2006), which, while useful, provides an incomplete and potentially biased view of genome-wide diversity. The availability of a full reference genome now enables deeper analyses of population structure, genome-wide variation, repeat content, and evolutionary history, greatly expanding the scope of research questions that can be addressed.

Notably, the identification and assembly of the W and Z sex chromosomes add critical value to this genomic resource. In many birds, sex chromosomes are hotspots for differentiation, harbor loci involved in reproductive isolation, and often exhibit different evolutionary dynamics than autosomes (Hooper et al., 2019; Luohao et al., 2019). Their inclusion facilitates investigations into sex-biased dispersal (Brom et al., 2018), differential gene expression (Mank & Ellegren, 2009), and the potential role of sex-linked loci in local adaptation (Nadachowska-Brzyska et al., 2019).

In addition to providing a valuable genomic resource for avian research, our findings have broad implications for conservation. The strong natal philopatry exhibited by *P. phaeopygia* (Cruz & Cruz, 1987) suggests that populations may be highly structured, potentially increasing their vulnerability to inbreeding and local extinction. With this genome, we will be able to genotype individuals across populations to assess gene flow, inbreeding levels, and fine-scale population differentiation, essential for defining conservation units and informing future management strategies.

Finally, the success of this project was made possible, in part, by sequencing DNA locally at the Galapagos Science Center (GSC). This approach ensured compliance with strict regulations governing the import and export of biological material. In addition, the use of portable ONT devices demonstrates the feasibility of conducting high-throughput sequencing in remote, resource-limited environments, while also contributing to the growing genomic research capacity in isolated regions such as the Galápagos Islands.

## Data Availability

Raw sequencing data is available on NCBI under BioProject SAMN49010279 and BioSample SAMN49010279 : GAPE_003. The genome assembly is deposited under the same BioProject on NCBI. All NCBI submissions will be made public at the time of publication. It can be accessed using this link (reviewers only): https://dataview.ncbi.nlm.nih.gov/object/PRJNA1275046?reviewer=thdo1oi1p8fetu460cr3lg7j7n

## Code Availability

Custom scripts are in the github.com/jaimechaves76/GalapaGenomes-Galapagos-Petrel-G3 github repository as well as the https://github.com/calacademy-research/assembly_etc github repository under the scripts folder. Files seq_summary_qscore_lens.sh for read statistics and asmstats.pl for assembly statistics are located there, as are script dependencies and basic_gff_stats.sh. Several scripts also use bioawk_cas from https://github.com/calacademy-research/bioawk.CAS, an enhanced version of Heng Li’s bioawk (https://github.com/lh3/bioawk).

## Acknowledgements

Special thanks to Jonathan Guillén for assistance in the field, the logistical support of the Galapagos Science Center, including Paúl Yepéz, Cristina Vintimilla, Gabriela Bautista, Jessenia Sotamba, and Sylvia Sotamba, and the Agencia de Regulación y Control de la Bioseguridad y Cuarentena para Galápagos (ABG), in particular, to Alberto Vélez. Athena Lam at the CAS Center for Comparative Genomics was instrumental in laboratory training. Work was done under the Permiso de Acceso al Recurso Genético-Ecuador MAATE-DBI-CM-2022-0249. This genome is part of the *GalapaGenomes* project funded by NSF (BRC-BIO) to J.A.C. under San Francisco State University IACUC A2021-27. This work is part of I.S.’s master’s thesis at San Francisco State University (SFSU).

This work was part of the ORG.one initiative (https://nanoporetech.com/oo).

## Author Contribution

J.A.C. and J.P.D. designed the project; J.A.C. acquired funding for the project; J.A.M. and G.P. conducted the lab work and produced all the genetic data; I.R.S. and J.B.H. performed the bioinformatic analysis; I.R.S. and J.B.H. wrote the manuscript with input from J.P.D., J.A.C., J.A.M., and V.F.; A.S. acquired the biological sample and funding for fieldwork. All authors checked and approved the final version of the manuscript.

